## Supplementary Material 1 for "Scientific modelling can be accessible, interoperable and user friendly: An example for pasture and livestock modelling"

### Glossary

General workflow differentiates the modules by colour (Fig. S1). This colour palette is applied when a model is run in a module other than its own. The workflows for each module follow the same legend (Fig. S2).

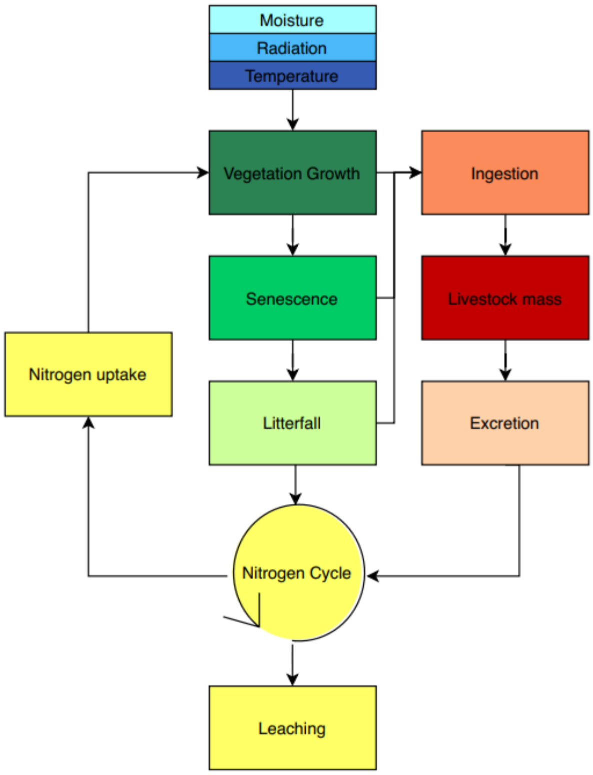


*Figure S 1 Module workflow*

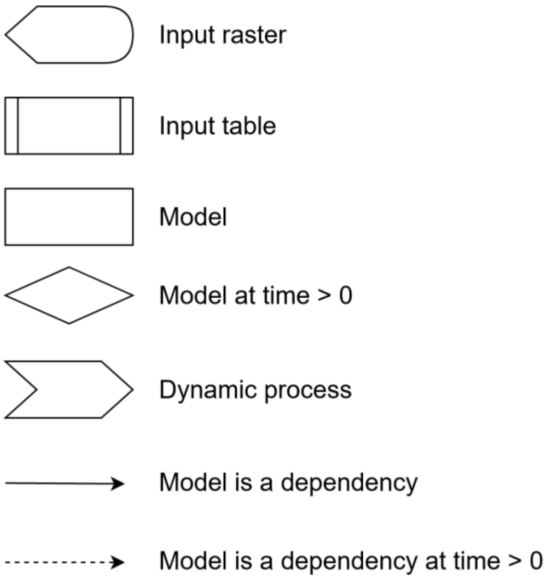


*Figure S 2 Model's data flow legend*

### 1. General Model

*Table S 1 General Models*

| Puerto id | Model | Description | Units |
|---|---|---|---|
| com | Type of Relevant Pastureland Plant | Primary plant species | |
| com2 | Type of Overstory Relevant Pastureland Plant | Secondary plant species | |
| n_c | Count of Cattle Individual | Number of animals per hectare | individuals/ha |
| n_m | Count of Mares Individual | Number of animals per hectare | individuals/ha |

### 2. Moisture

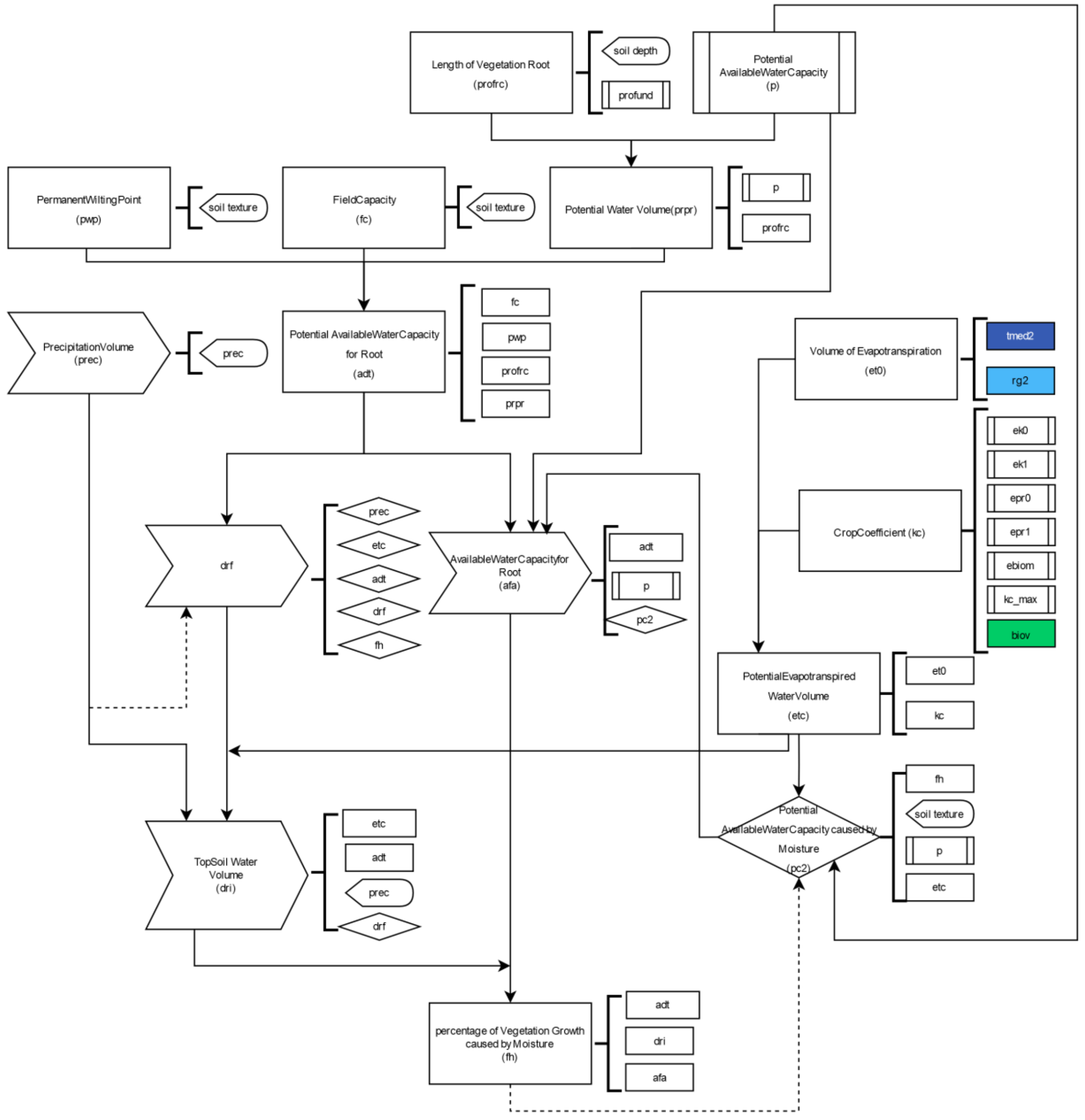


*Figure S 3 Workflow of the moisture module*

*Table S 2 Moisture models*

| Puerto id | Model | Description | Units |
|---|---|---|---|
| adt | Potential AvailableWaterCapacity for Root | Potential available soil water in the root zone | mm |
| afa | AvailableWaterCapacity for Root | The readily available soil water in the root zone | mm |
| drf | | Initial depletion | |
| dri | TopSoil Water Volume | Volume of water in the root zone caused by the limits on root zone depletion by evapotranspiration | mm |
| ebiom | | Biomass when Kc (crop coefficient) is 1 | $g/m^2$ |
| ek0 | | Minimum and constant Crop Coefficient | |
| ek1 | | Definition of the function slope between Kc (Crop coefficient) and biov (living biomass) | |
| epr0 | | Proportion of the biov (living biomass) reference below which Kc (Crop Coefficient) is minimum and constant (ek0) | |
| epr1 | | Proportion of the biov (living biomass) reference at which kc (Crop Coefficient) = kcmax (Maximum Crop coefficient) | |
| et0 | Volume of Evapotranspiration | Reference evapotranspiration | mm |
| etc | PotentialEvapotranspiredWaterVolume | Potential Evapotranspiration | mm |
| fc | FieldCapacity | Field Capacity | $m^3/m^3$ |
| prec | PrecipitationVolume | Volume of precipitation | mm |
| fh | Percentage of VegetationGrowth caused by Moisture | Coefficient between 0 and 1 that calculates vegetation water stress through soil moisture and vegetation characteristics | [0-1] |
| kc | CropCoefficient | Crop coefficient | |
| kcmax | | Maximum value of crop coefficient following rain or irrigation | |
| p | Potential AvailableWaterCapacity | Average fraction of total available soil water that can be depleted from the root zone before moisture stress (reduction in ET) occurs | [0-1] |
| pc2 | Potential AvailableWaterCapacity caused by Moisture | Adjustment of p (AvailableWaterCapacity) for different etc (PotentialEvapotranspiredWaterVolume) | |
| soil_depth | SoilDepth | Soil depth | mm |
| prec | PrecipitationVolume | Precipitation | mm |
| profr | Maximum Length of Root | Root depth when there are no bedrock limitations | mm |
| profrc | Length of Vegetation Root | Root depth considering limitations | mm |
| prpr | Potential Water Volume | Proportion of soil water content available to roots | |
| pwp | PermanentWiltingPoint | Permanent wilting point, defined as the minimal amount of water in the soil that the plant requires to avoid wilting | $m^3/m^3$ |
| soil_texture | Type of SoilTexture | Soil texture | |

3. Radiation

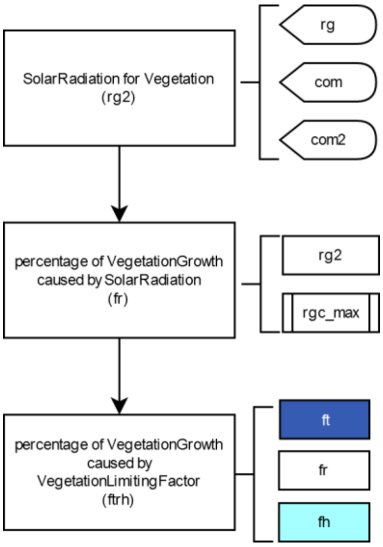


*Figure S 4 Workflow of the moisture module*

*Table S 3 Radiation models*

| Puerto id | Model | Description | Units |
|---|---|---|---|
| fr | Percentage of Vegetation Growth caused by SolarRadiation | Percentage of radiation that limits vegetation growth. 0: No growth (maximum limitation), 1: Maximum growth (no limitation) | [0-1] |
| ftrh | Percentage of Vegetation Growth caused by VegetationLimitingFactor | Set of climatic factors (temperature, radiation and soil moisture) limiting vegetation growth | [0-1] |
| rg | SolarRadiation | Incidence of solar radiation | MJ |
| rg2 | SolarRadiation for Vegetation | Incidence of solar radiation over vegetation | $MJ/m^2$ |
| rgc_max | Maximum PhotosyntheticallyActiveRadiation | Incidence of solar radiation for maximum photosynthetic capacity | $MJ/m^2$ |

4. Temperature

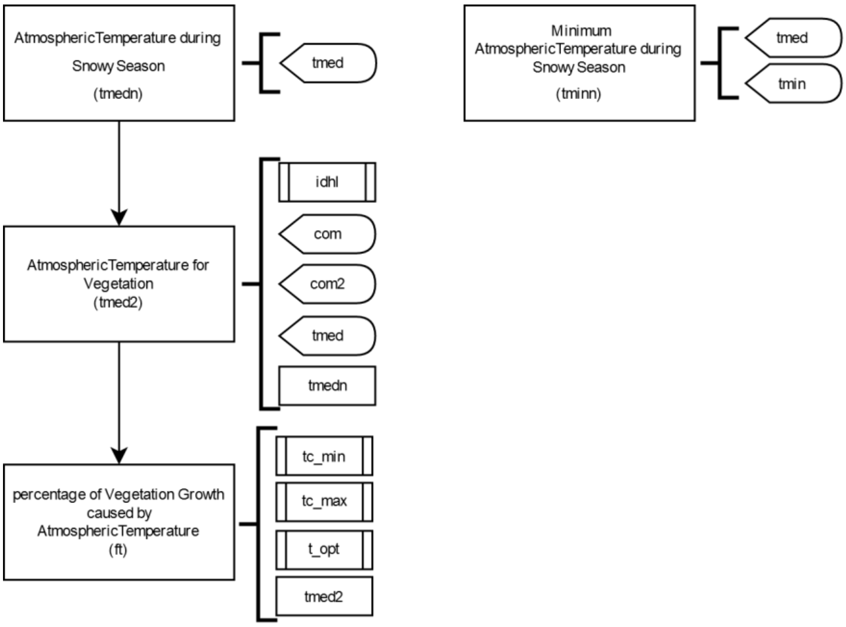


*Figure S 5 Workflow of the temperature module*

*Table S 4 Temperature models*

| Puerto id | Model | Description | Units |
|---|---|---|---|
| ft | Percentage of Vegetation Growth caused by AtmosphericTemperature | Percentage of Atmospheric temperature that limits vegetation growth. 0: No growth (maximum limitation), 1: Maximum growth (no limitation) | [0-1] |
| idhl | Value of VegetationStratum | Vertical vegetation stratum coded as 1: ground layer; 2: shrubs layer; 3: midstory layer; 4: canopy layer | |
| tc_max | Maximum AtmosphericTemperature causing Vegetation Growth | Maximum temperature limiting vegetation growth | °C |
| tc_min | Minimum AtmosphericTemperature causing Vegetation Growth | Minimum temperature limiting vegetation growth | °C |
| tmed | AtmosphericTemperature | Mean atmospheric temperature | °C |
| tmed2 | AtmosphericTemperature for Vegetation in Celsius | Vegetation temperature under woody plants | °C |
| tmedn | AtmosphericTemperature during Snowy Season | Mean atmospheric temperature over vegetation in snowy conditions | °C |
| tmin | Minimum AtmosphericTemperature | Minimum atmospheric temperature | °C |
| tminn | Minimum AtmosphericTemperature during Snowy Season in Celsius | Minimum atmospheric temperature over vegetation in snowy conditions | °C |
| topt | AtmosphericTemperature causing Maximum Vegetation Growth | Optimum temperature for vegetation growth | °C |

5. Vegetation growth

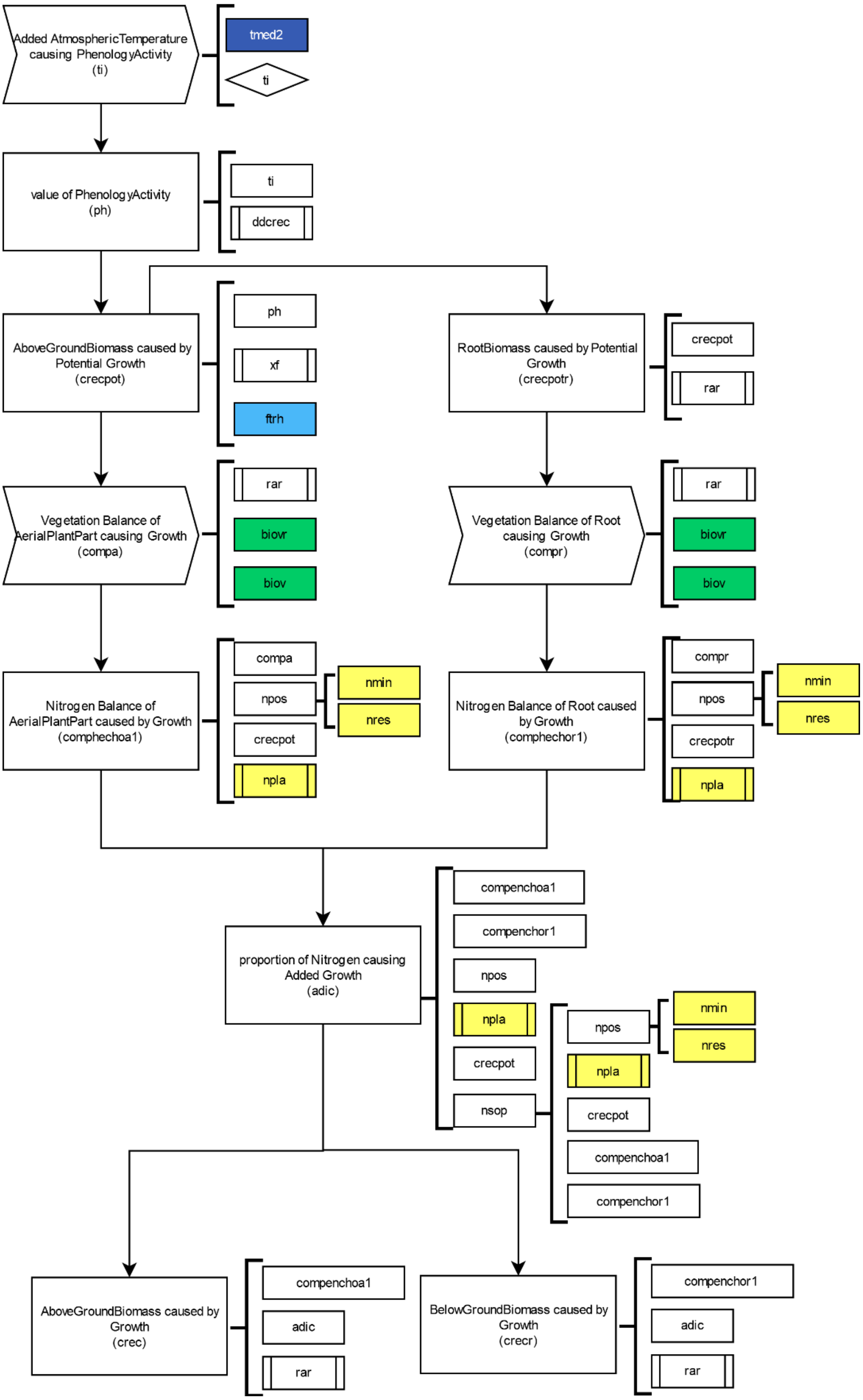


*Figure S 6Workflow of the vegetation growth module*

*Table S 5 Vegetation growth models*

| Puerto id | Model | Description | Units |
|---|---|---|---|
| adic | Proportion of Nitrogen causing Added Growth | Proportion of the remainder nitrogen causing extra vegetation growth | [0-1] |
| comp | | Function to compute compa and compr | |
| compa | Vegetation Balance of AerialPlantPart causing Growth | The remainder of net primary production is used for aerial vegetation growth. | |
| comphechoa | | Function to calculate comphechoa1 | |
| comphechoa1 | Nitrogen Balance of AerialPlantPart caused by Growth | Normalised concentration of nitrogen in the aerial part of the plant | [0-1] |
| comphechor | | Function to calculate comphechoar1 | |
| comphechor1 | Nitrogen Balance of Root caused by Growth | Normalised concentration of nitrogen in the root part of the plant | [0-1] |
| compr | Vegetation Balance of Root causing Growth | The remainder of net primary production used for root growth | |
| crec | AboveGroundBiomass caused by Growth | Actual growth of above ground biomass | $g/m^2$ |
| crecpot | Potential AboveGroundBiomass caused by Potential Growth | Potential growth of above ground biomass | $g/m^2$ |
| crecpotr | Potential RootBiomass caused by Potential Growth | Potential growth of below ground biomass | $g/m^2$ |
| crecr | RootBiomass caused by Growth | Actual growth of below ground biomass | $g/m^2$ |
| ddcrec | Added AtmosphericTemperature causing Active Growth in Celsius | Growing degree days (GDD) accumulated, a commonly used measure of thermal accumulation | °C*day |
| npos | | Function that sums nres (nitrogen causing vegetation growth) and mnim (inorganic nitrogen) | $g/m^2$ |
| ph | Occurrence of PhenologyActivity | Phenological state of vegetation | [0-1] |
| rar | Proportion of RootBiomass in Balance Vegetation | Relation between the growth of aerial and root system | [0-1] |
| ti | Added AtmosphericTemperature causing PhenologyActivity in Celsius | Accumulated temperature necessary to complete a phenological state | °C |
| xf | Maximum Biomass caused by Growth | Theoretical vegetation growth without limitations | $g/m^2$*day |

### 6. Senescence

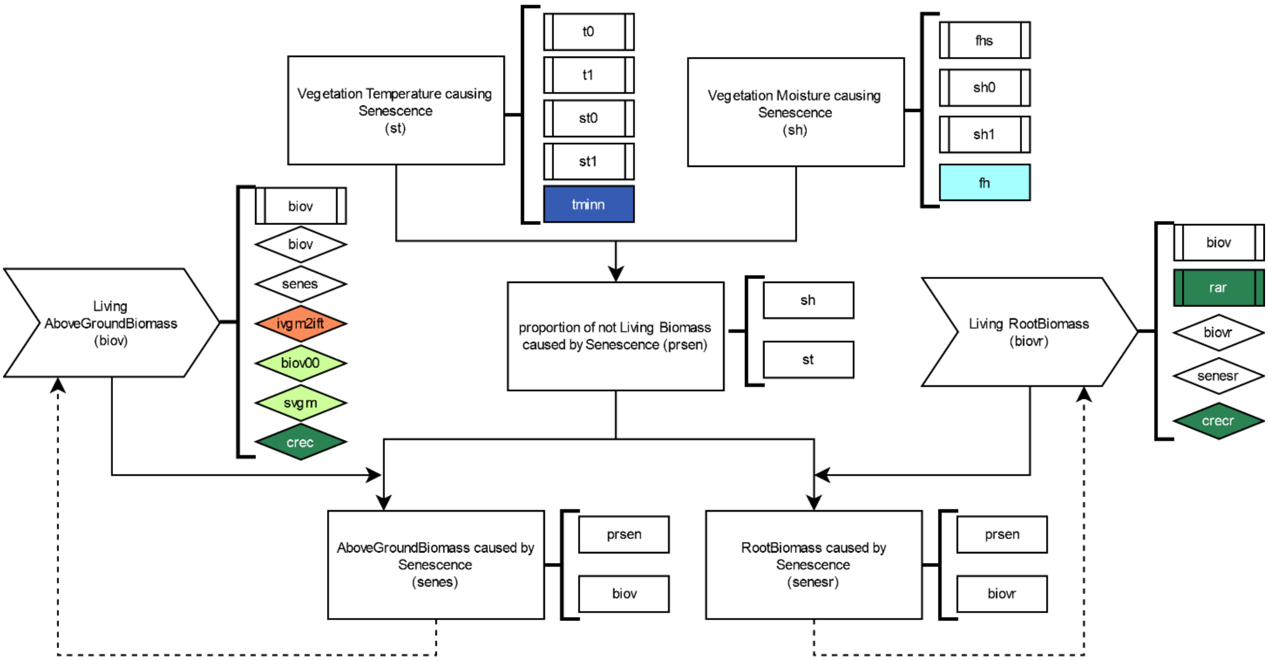


*Figure S 7 Workflow of the senescence module*

*Table S 6 Senescence models*

| Puerto id | Model | Description | Unit |
|---|---|---|---|
| biov | Living AboveGroundBiomass | Living above ground biomass | $g/m^2$ |
| biovr | Living RootBiomass | Living below ground biomass | $g/m^2$ |
| fhs | Proportion of Moisture causing Minimum Senescence | Level of moisture (fh) with minimum senescence | |
| prsen | Proportion of not Living Biomass caused by Senescence | Maximum proportion of senescence caused by temperature or moisture | [0-1] |
| senes | AboveGroundBiomass caused by Senescence | Amount of above ground biomass dying caused by senescence | $g/m^2$ |
| senesr | RootBiomass caused by Senescence | Amount of below ground biomass dying caused by senescence | $g/m^2$ |
| sh | Vegetation Moisture causing Senescence | Proportion of leave senescence caused by moisture | [0-1] |
| sh0 | | Maximum proportion of leaves dying due to moisture | |
| sh1 | | Proportion of minimum leave senescence caused by soil moisture | |
| st | Vegetation AtmosphericTemperature causing Senescence | Proportion of senescence caused by temperature | [0-1] |
| st0 | | Maximum proportion of leaves dying due to temperature | |
| st1 | | Temperature causing minimal senescence | |
| t0 | AtmosphericTemperature causing Maximum Senescence | Atmospheric temperature producing maximum senescence | °C |
| t1 | AtmosphericTemperature causing Minimum Senescence | Atmospheric temperature producing minimum senescence | °C |

7. Litterfall

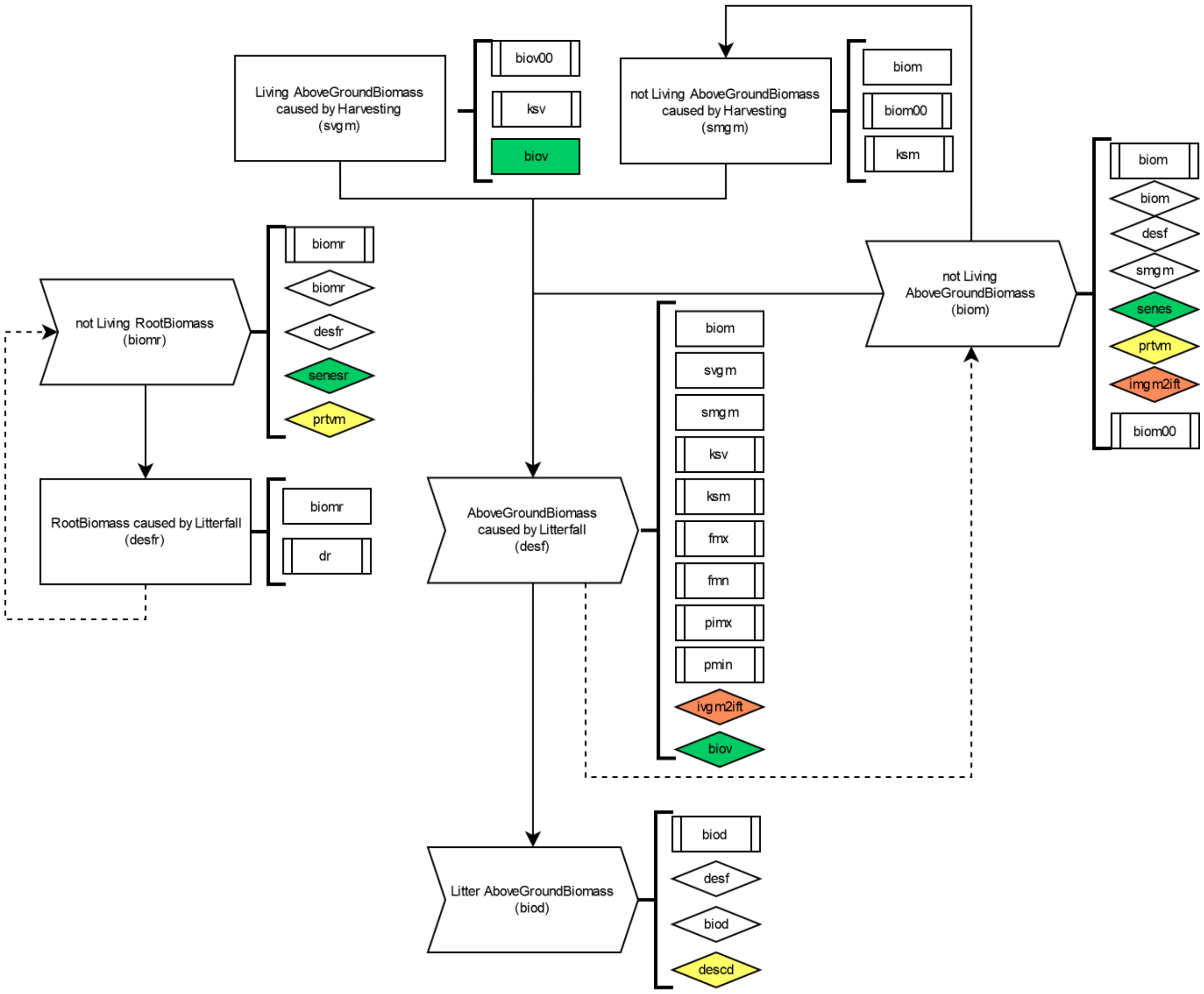


*Figure S 8 Workflow of the litterfall module*

*Table S 7 Litterfall models*

| Puerto id | Model | Description | Unit |
|---|---|---|---|
| biod | Litter AboveGroundBiomass | Amount of litterfall related to above ground biomass | $g/m^2$ |
| biom | Not Living AboveGroundBiomass | Dead standing above ground biomass | $g/m^2$ |
| biom00 | | Standing dead biomass remaining after harvest | $g/m^2$ |
| biomr | Not Living RootBiomass | Dead standing root biomass | $g/m^2$ |
| biov00 | | Living standing biomass remaining after harvest | $g/m^2$ |
| desf | AboveGroundBiomass caused by Litterfall | Process of litterfall related to above ground biomass | $g/m^2$ |
| desfr | RootBiomass caused by Litterfall | Process of litterfall related to root biomass | $g/m^2$ |
| dr | | Rate of root litterfall | |
| fmn | Proportion of Minimum Litterfall in not Living AboveGroundBiomass | Minimum proportion of litterfall caused by livestock | day |
| fmx | Proportion of Maximum Litterfall in not Living AboveGroundBiomass | Maximum proportion of litterfall caused by livestock | day |
| ksm | Proportion of not Living AboveGroundBiomass in Harvesting | Harvesting efficiency of dead above ground biomass | |
| ksv | Proportion of Living AboveGroundBiomass in Harvesting | Harvesting efficiency of living above ground biomass | |

| | | | |
|---|---|---|---|
| pimn | Proportion of AboveGroundBiomass in Minimum Litterfall | Proportion of ingested biomass by livestock with minimum fall | day |
| pimx | Proportion of AboveGroundBiomass in Maximum Litterfall | Proportion of ingested biomass by livestock with maximum fall | day |
| smgm | Not Living AboveGroundBiomass caused by Harvesting | Amount of dead biomass harvested | $g/m^2$ |
| svgm | Living AboveGroundBiomass caused by Harvesting | Amount of living biomass harvested | $g/m^2$ |

8. Livestock Ingestion

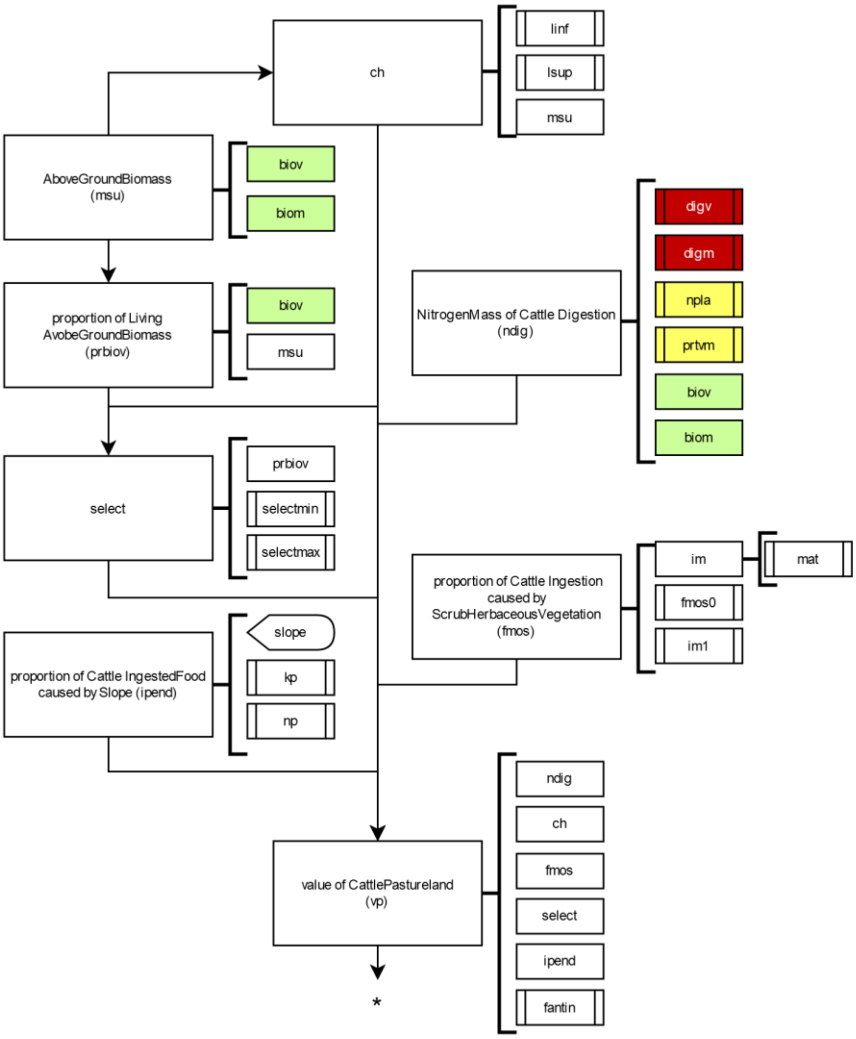

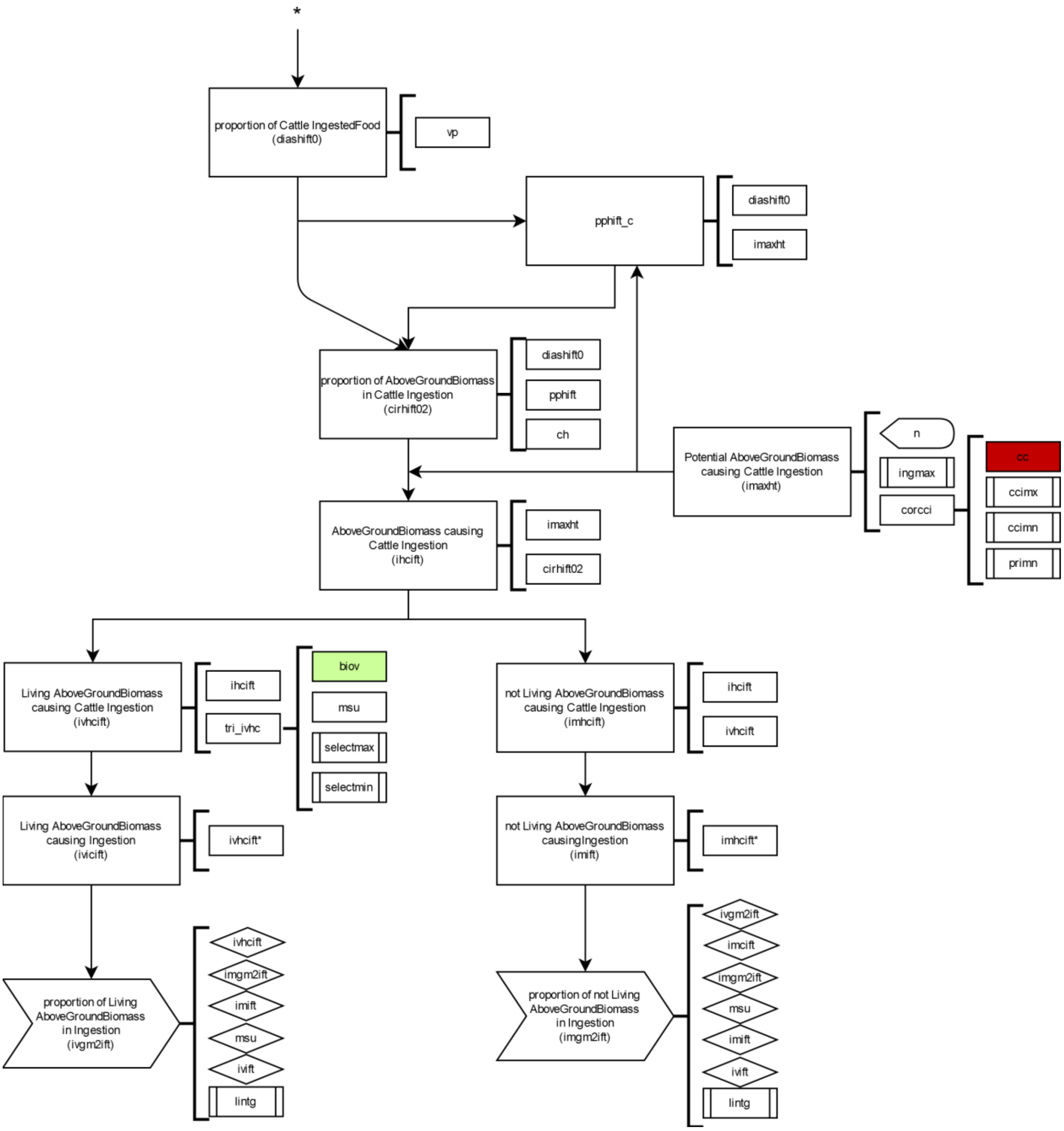


*Figure S 9 Workflow of the ingestion module*

*Table S 8 Ingestion models*

| Puerto id | Model | Description | Units |
|---|---|---|---|
| ccimn | | Body condition at minimal potential intake | |
| ccimx | | Body condition at maximum potential intake | |
| ch | | Coefficient (01) of regulation of livestock intake according to available biomass | [0-1] |
| cirhift01 | | Potential of total intake proportion corresponding to type of grass | |
| cirhift02 | Proportion of AboveGroundBiomass in Cattle Ingestion | Real proportion of the herd's ingestion corresponding to type of grass | |
| corcci | | Function to calculate the livestock body condition | |

| | | | |
|---|---|---|---|
| diashift0 | Proportion of Cattle IngestedFood | Estimates the proportion of grazing if there were no overlap with others' herds | |
| fantin | | Maximum proportion of (01) a type of grass can be part of the animal's daily diet | [0-1] |
| fmos | Proportion of Cattle Ingestion caused by ScrubHerbaceousVegetation | Difficulty of livestock access due to excess shrub growth (0: not accessible; 1 : fully accessible) | [0-1] |
| fmos0 | | Minimum accessibility of livestock | |
| ihcift | AboveGroundBiomass causing Cattle Ingestion | Amount of above ground biomass ingested by each type of livestock | |
| im | Proportion of ScrubHerbaceousVegetation causing Cattle Movement | Level of livestock accessibility due to the proportion of shrubs (0: no shrubs, 1: maximum proportion of shrubs) | |
| im0 | | Level of shrub intake with minimal livestock accessibility | |
| im1 | | Level of shrub intake with maximal livestock accessibility | |
| imaxht | Potential AboveGroundBiomass causing Cattle Ingestion | Potential ingestion without biomass limitation | |
| imgm2ift | Proportion of not Living AboveGroundBiomass in Ingestion | Proportion of ingested nonliving biomass in the whole of the ingested biomass | |
| imhcift | Not Living AboveGroundBiomass causing Livestock Ingestion | Nonliving above ground biomass ingested by each type of livestock | |
| imift | Not Living AboveGroundBiomass causing Ingestion | Nonliving above ground biomass ingested by livestock | |
| ingmax | | Maximum ingestion | kg/day |
| ipend | Proportion of Cattle IngestedFood caused by Slope | Coefficient (01) of grazing capacity according to slope | [0-1] |
| ivgm2ift | Proportion of Living AboveGroundBiomass in Ingestion | Proportion of ingested living biomass out of all ingested biomass | |
| ivhcift | Living AboveGroundBiomass causing Cattle Ingestion named ivhcift | Living above ground biomass ingested by each type of livestock | |
| ivift | Living AboveGroundBiomass causing Ingestion | Living above ground biomass ingested by livestock | |
| kp | | Parameter that indicates the slope when the function of the model "proportion of Cattle IngestedFood caused by Slope" is 0.5 | % |
| linf | | Amount of aerial biomass that prevents grazing | |
| lintg | | Amount of biomass below which there is no intake | g/m2 |
| lsup | | Amount of aerial biomass that allows grazing | |
| mat | | Probability of accessibility and livestock movement through vegetation | [0-1] |
| msu | AboveGroundBiomass | Standing above ground biomass (living and nonliving) | g/m^2 |
| ndig | Nitrogen Mass of Cattle Digestion | Digestible nitrogen mass of standing vegetation | g/m^2 |
| np | | Parameter indicating the decrease in the "proportion of feed intake by livestock caused by the slope" model when the slope increases | |
| pphift | | Potential of total grazing proportion of a given pasture corresponding to a herd | |
| prbiov | Proportion of Living AvobeGroundBiomass | Proportion of living above ground biomass | |

| | | | |
|---|---|---|---|
| primn | | Proportion of potential intake to “ccimn” model | |
| select | | Coefficient (01) indicating the livestock ability to select the living against the dead part of the vegetation | [0-1] |
| selectmax | Maximum proportion of Living AboveGroundBiomass in IngestedFood | Proportion of living biomass that allows the livestock to select only living vegetation | |
| selectmin | Minimum proportion of Living AboveGroundBiomass in IngestedFood | Proportion of living biomass that prevents the livestock from selecting living vegetation | |
| slope | Slope | Terrain inclination | % |
| tri_ivhc | | Function to calculate ingestion model | |
| vp | Value of Cattle Pastureland | Dimensionless index allowing estimation of pasture quality according to its composition | gN/m^2 |

### 9. Livestock Excretion

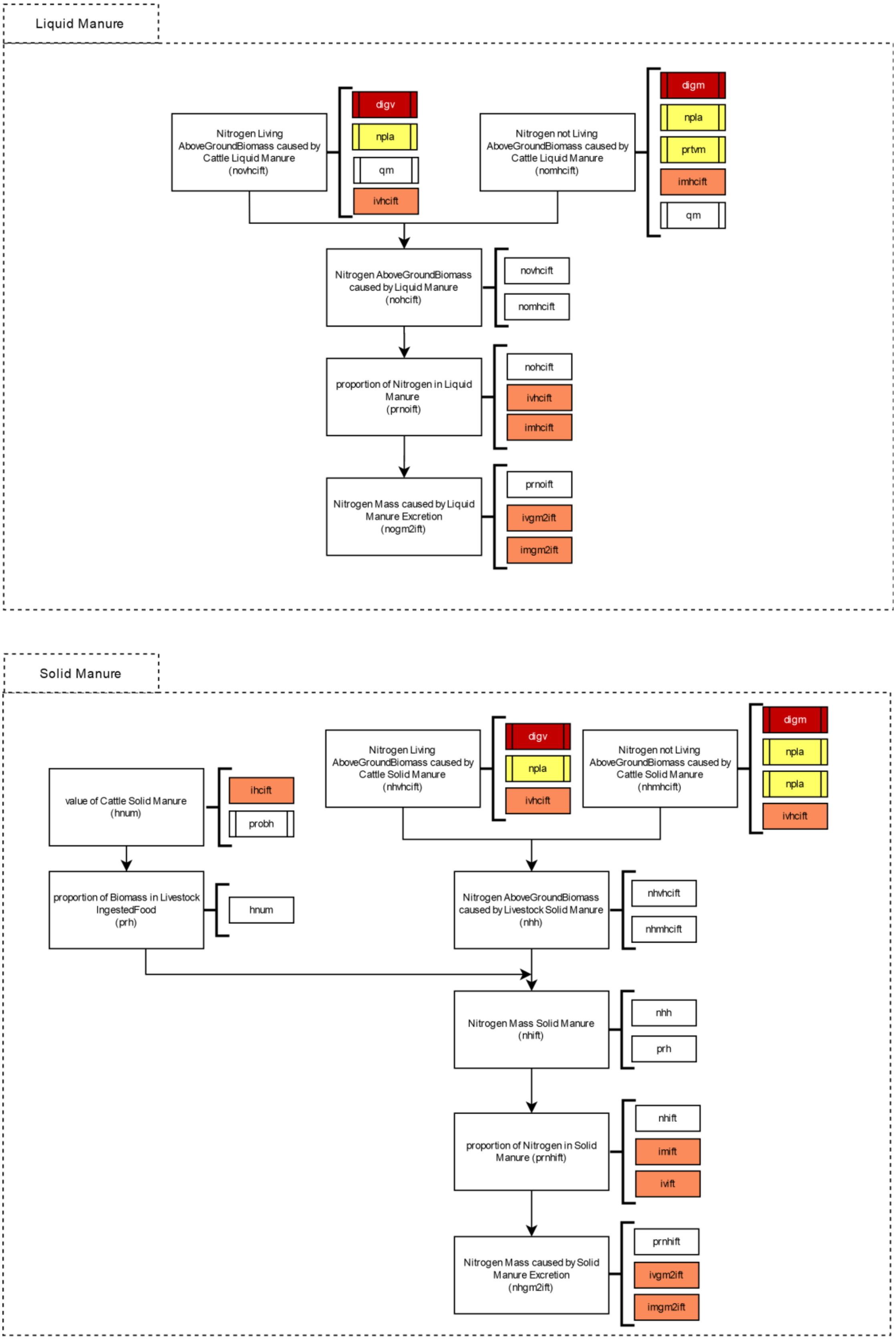


*Figure S 10 9Workflow of the excretion module*

*Table S 9 Excretion models*

| Puerto id | Model | Description | Units |
|---|---|---|---|
| qm | | Coefficient of energy metabolism | |
| nhmhcift | Nitrogen not Living AboveGroundBiomass caused by Cattle Solid Manure | Nitrogen content of nonliving AboveGroundBiomass caused by Cattle Solid Manure | |
| hnum | Cattle Solid Manure | Livestock Solid Manure | |
| nhgm2ift | Nitrogen Mass caused by Solid Manure Excretion | Spatial distribution of solid manure nitrogen concentration | |
| nhhift | Nitrogen AboveGroundBiomass caused by Livestock Solid Manure | Nitrogen AboveGroundBiomass caused by Livestock Solid Manure | |
| nhift | Nitrogen Mass Solid Manure | Nitrogen concentration of solid manure | |
| nhvhcift | Nitrogen Living AboveGroundBiomass caused by Cattle Solid Manure | Nitrogen mass of living biomass in manure | |
| nogm2ift | Nitrogen Mass caused by Liquid Manure Excretion | Concentration of nitrogen in liquid manure | |
| nohcift | Nitrogen AboveGroundBiomass caused by Liquid Manure | Nitrogen concentration of liquid manure caused by total of aboveground biomass ingestion | |
| nomhcift | Nitrogen not Living AboveGroundBiomass caused by Cattle Liquid Manure | Nitrogen concentration of liquid manure caused by total of non living aboveground biomass ingestion | |
| norinmhif | Nitrogen not Living AboveGroundBiomass caused by Cattle Liquid Manure | Nitrogen concentration of liquid manure caused by non living aboveground biomass ingestion | |
| novhcift | Nitrogen Living AboveGroundBiomass caused by Cattle Liquid Manure | Nitrogen concentration of liquid manure caused by aboveground biomass ingestion | |
| prh | Proportion of Biomass in Livestock IngestedFood | proportion of Biomass in Livestock IngestedFood | |
| prnhift | Proportion of Nitrogen in Solid Manure | Proportion of nitrogen in manure | |
| prnoift | Proportion of Nitrogen in Liquid Manure | Proportion of nitrogen in liquid part of manure | |
| probh | Occurrence of Excretion | Probability of manure left in a plant community compared to its use | [0-1] |

10. Livestock mass

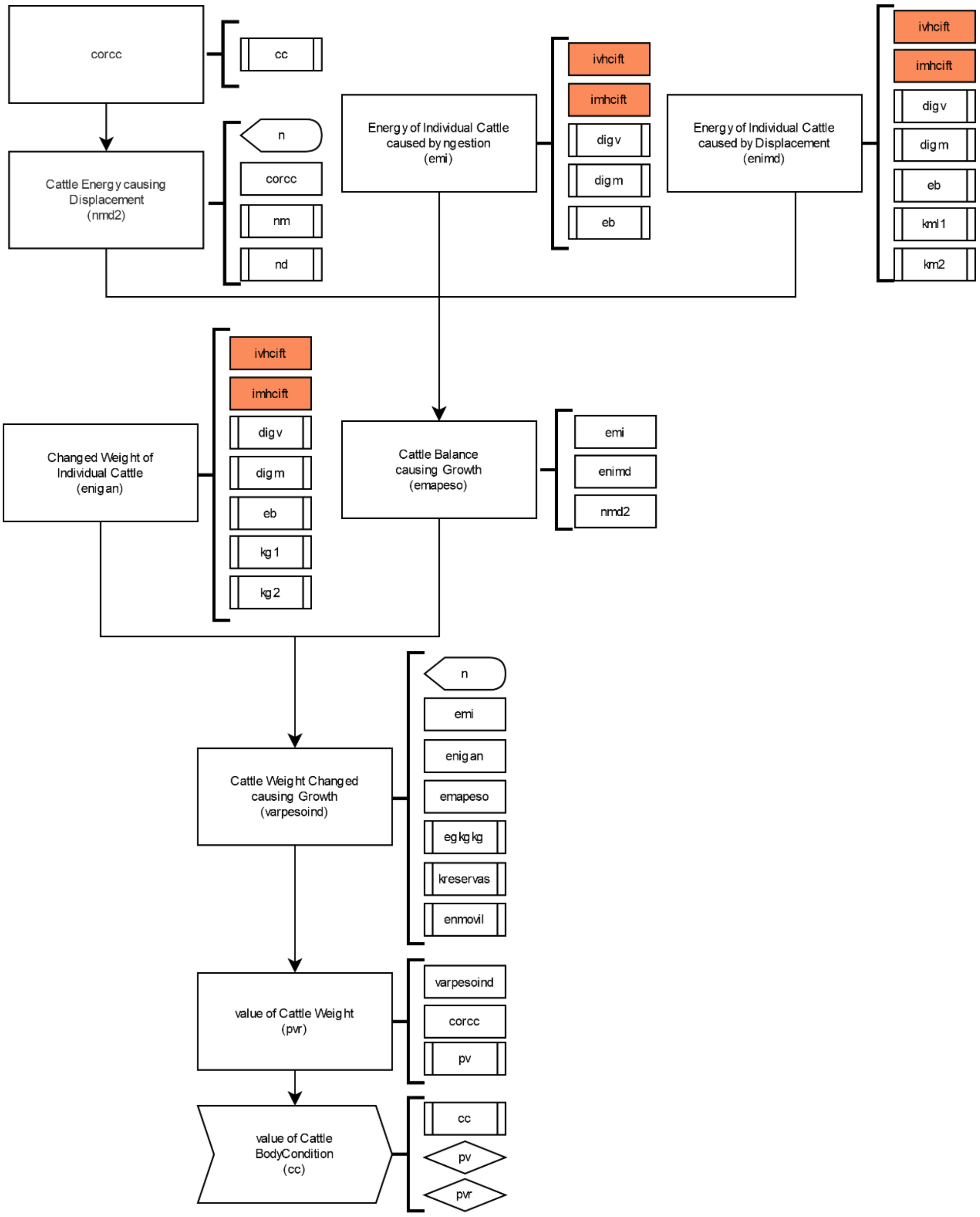


*Figure S 11 Workflow of the livestock mass module*

*Table S 10 Livestock mass models*

| Puerto id | Model | Description | Units |
|---|---|---|---|
| cc | Value of Cattle BodyCondition | Body Condition | [0-5] |
| digm | Proportion of not Living AboveGroundBiomass in Cattle Digestion | Digestibility of nonliving aerial biomass | |
| digv | Proportion of Living AboveGroundBiomass in Cattle Digestion | Digestibility of living aerial biomass | |

| | | | |
|---|---|---|---|
| eb | | Gross energy value | MJ/kg |
| egkg | | Energy needed to increase 1 kg of body weight | MJ/kg |
| emapeso | Cattle Balance causing Growth | Balance between ingested energy and needs | |
| emi | Energy of Individual Cattle caused by Ingestion | Metabolisation of ingested energy in livestock | MJ/t |
| enigan | Changed Weight of Individual Cattle | Ingested energy that implies change of weight | |
| enimd | Energy of Individual Cattle caused by Displacement | Ingested energy that implies movement and displacement | |
| enmovil | | Energy value of mobilizing body reserves | MJ/kg |
| kg1 | | Energy efficiency to gain weight | |
| kg2 | | Metabolism energy efficiency for gain weight | |
| km2 | | Metabolism energy efficiency for maintenance and mobility | |
| kml1 | | Efficiency of energy use for lactation | |
| kreservas | | Efficiency of metabolizable energy used for food supplies mobilization | |
| nmd2 | Cattle Energy causing Displacement | Livestock movement and displacement energy | MJ/d |
| pv | | Reference live weight for each type of livestock | kg |
| pvr | Value of Cattle Weight | Reference body weight | |
| varpesoind | Cattle Weight Changed causing Growth | Change of weight | |

### 11. Nitrogen cycle

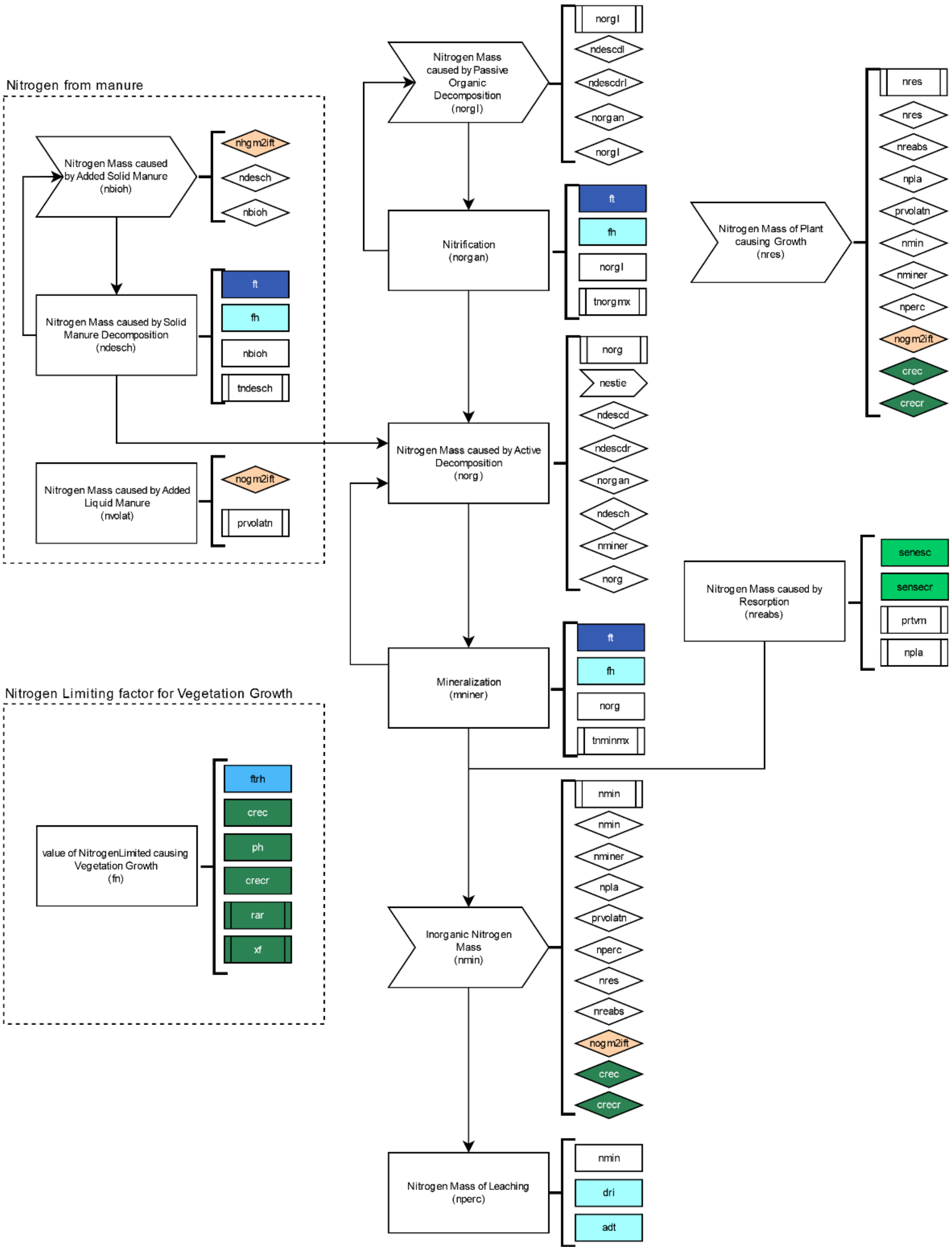


*Figure S 12 First part of the workflow of the nitrogen module*

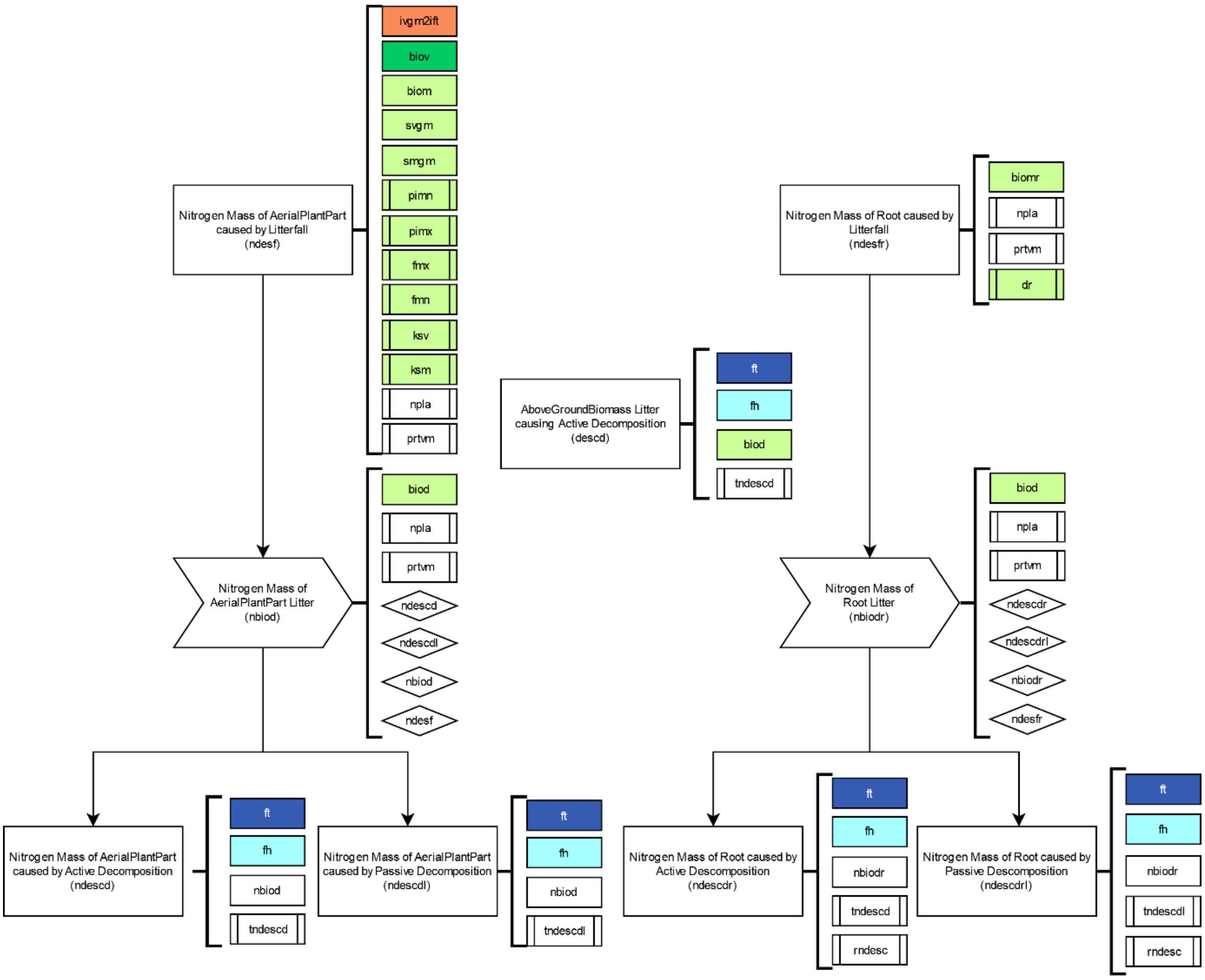


*Figure S 13 Second part of the workflow of the nitrogen module*

*Table S 11 Nitrogen models*

| Puerto id | Model | Description | Units |
|---|---|---|---|
| descd | AboveGroundBiomass Litter causing Active Decomposition | Amount of litterfall of aerial vegetation biomass in active decomposition process | $g/m^2$ |
| descdl | AboveGroundBiomass Litter causing Passive Decomposition | Amount of litterfall of aerial vegetation biomass in passive decomposition process | $g/m^2$ |
| fn | Value of NitrogenLimited causing Vegetation Growth | Range of nitrogen mass that limits vegetation growth | [0-1] |
| nbiod | Nitrogen Mass of AerialPlantPart Litter | Amount of nitrogen in litterfall from aerial vegetation part | $g/m^2$ |
| nbiodr | Nitrogen Mass of Root Litter | Amount of nitrogen in litterfall from root vegetation part | $g/m^2$ |
| nbioh | Nitrogen Mass caused by Added Solid Manure | Amount of nitrogen accumulation from solid manure | $g/m^2$ |
| ndescd | Nitrogen Mass of AerialPlantPart caused by Active Decomposition | Nitrogen mass in active decomposition (aerial vegetation part) | $g/m^2$ |
| ndescdl | Nitrogen Mass of AerialPlantPart caused by Passive Decomposition | Nitrogen mass in passive decomposition (aerial vegetation part) | $g/m^2$ |
| ndescdr | Nitrogen Mass of Root caused by Active Decomposition | Nitrogen mass in active decomposition (root vegetation part) | $g/m^2$ |
| ndescdrl | Nitrogen Mass of Root caused by Passive Decomposition | Nitrogen mass in passive decomposition (root vegetation part) | $g/m^2$ |

| | | | |
|---|---|---|---|
| ndesch | Nitrogen Mass caused by Solid Manure Decomposition | The amount of nitrogen that changes from solid manure decomposition to active organic nitrogen | $g/m^2$ |
| ndesf | Nitrogen Mass of AerialPlantPart caused by Litterfall | Amount of nitrogen in aerial biomass litterfall | $g/m^2$ |
| ndesfr | Nitrogen Mass of Root caused by Litterfall | Amount of nitrogen in root biomass litterfall | $g/m^2$ |
| nestie | Nitrogen Mass causing Fertilization | Amount of organic nitrogen added by farmer as fertilizer | $g/m^2$ |
| nmin | Inorganic Nitrogen Mass | Amount of inorganic nitrogen | $g/m^2$ |
| nminer | Mineralization | Mineralization process | $g/m^2$ |
| norg | Nitrogen Mass caused by Active Decomposition | Amount of nitrogen in active decomposition | $g/m^2$ |
| norgan | Nitrification | Nitrification process | |
| norgl | Nitrogen Mass caused by Passive Organic Decomposition | Amount of nitrogen in passive organic decomposition | $g/m^2$ |
| nperc | Nitrogen Mass of Leaching | Amount of leaching nitrogen | $g/m^2$ |
| npla | Proportion of Nitrogen in Living AboveGroundBiomass | Proportion of nitrogen in living above ground biomass | |
| nreabs | Nitrogen Mass caused by Resorption | Amount of nitrogen in resorption process | $g/m^2$ |
| nres | Nitrogen Mass of Plant causing Growth | Amount of nitrogen causing vegetation growth | $g/m^2$ |
| nvolat | Nitrogen Mass caused by Added Liquid Manure | Amount of nitrogen accumulation from liquid manure | $g/m^2$ |
| prtvm | Proportion of Nitrogen Mass in Resorption | Proportion of nitrogen in resorption process | |
| prvolatn | Proportion of Nitrogen in Volatilization | Proportion of volatilized liquid manure | |
| tndescd | | Maximum proportion of litterfall nitrogen in active organic matter | |
| tndescdl | | Maximum proportion of litterfall nitrogen in passive organic matter | day |
| tndesch | | Maximum proportion of nitrogen from solid manure to soil | day |
| tnminmx | | Maximum proportion of organic nitrogen in mineralization process | day |
| tnorgmx | | Maximum proportion of nitrogen from passive to active organic matter | day |
